# Structured illumination to spatially map chromatin motions

**DOI:** 10.1101/191098

**Authors:** Keith Bonin, Amanda Smelser, Naike Salvador Moreno, George Holzwarth, Kevin Wang, Preston Levi, Pierre-Alexandre Vidi

## Abstract

We describe a simple optical method that creates structured illumination of a photoactivatable probe and apply this method to characterize chromatin motions in the nuclei of live cells. A laser beam coupled to a diffractive optical element at the back focal plane of an excitation objective generates an array of near diffraction-limited beamlets with FWHM of 340±30 nm, which simultaneously photoactivate a 7x7 matrix pattern of GFP-labeled histones, with spots 1.70 μm apart. From the movements of the photoactivated spots, we map chromatin diffusion coefficients at multiple microdomains of the cell nucleus. The results show correlated motions of nearest chromatin microdomain neighbors, whereas chromatin movements are uncorrelated at the global scale of the nucleus. The method also reveals DNA damage-dependent decrease in chromatin diffusion. The DOE instrumentation can easily and cheaply be implemented on commercial inverted fluorescence microscopes to analyze adherent cell culture models. A protocol to measure chromatin motions in non-adherent human hematopoietic stem and progenitor cells is also described. We anticipate that the method will contribute to the identification of the mechanisms regulating chromatin mobility, which influences most genomic processes and may underlie the biogenesis of genomic translocations associated with hematologic malignancies.

## 1 Introduction

Dynamic motions of chromatin are thought to critically influence genomic processes such as gene expression, DNA replication, DNA repair, and the biogenesis of genomic translocations. Chromatin motions follow stochastic constrained random walks (1), which complicates the interpretation of their biological significance. Most studies on chromatin mobility rely on tracking artificial DNA arrays integrated in the genome (2-4), or more recently, DNA repeats by CRISPR/dCas9 imaging (5, 6). These tracking approaches have several limitations. First, artificial DNA arrays do not fully reproduce the complex chromatin organization (7), and chromatin binding of the bulky dCas9-GFP reporter may generate a drag affecting chromatin dynamics. Second, only a few measurements per cell are typically possible, from which a global assessment of chromatin dynamics is difficult. As such, new optical techniques are needed to interrogate a large number of regions in the cell nucleus, over meaningful periods of time. Photoactivatable histone probes have been used as an alternative to study chromatin motions in a near-native chromatin context (8-10). With this approach, the optical method used for photoactivation is critical, and conventional scanning confocal microscopy commonly used for analysis does not allow to simultaneously illuminate and photoactivate multiple sub-cellular regions.

Structured illumination has been increasingly applied over the last decades to improve performance in wide-field microscopy (11, 12). In addition to superresolution microscopy, applications for structured illumination include optical trapping, surface profiling, quantitative phase imaging in biological systems, and optical sectioning achieved by illumination with incoherent light (11, 13, 14). Here, we apply structured illumination for simultaneous photoactivation of chromatin reporters throughout the nucleus of live cells. Our structured light pattern is produced with a diffractive optical element (DOE) module, implemented as a simple modification to a commercial inverted microscope system. By enabling parallel measurements in native chromatin environments, the approach circumvents the limitations discussed above. It yields robust values of diffusion at multiple points within a single cell, thereby reducing the need for agglomerating population-based measurements, and provides spatial information on chromatin motions.

## 2 Materials and Methods

### 2.1 Instrument design

The optical setup is illustrated in Fig. 1. The laser source is a 30 mW fiber pigtailed diode laser (Thorlabs LP405-SF30) producing light at a wavelength of 405 nm, the photoactivation wavelength of photoactivatible green florescent protein (PAGFP). The fiber is single mode to produce a clean Gaussian spatial profile that is collimated by a metal mirror (Thorlabs RC12FC-F01), oriented to produce a reflected beam at 90° to the incoming axis, and designed to produce a beam of 12 mm in diameter. The collimated beam reflected downward by the collimating mirror is transmitted through a fused quartz diffractive optical element (DOE – Holo/Or MS-571-S-Y-X) designed to produce a 7x7 pattern of near diffraction-limited spots with high efficiency (~76%) at 405 nm. The DOE has a clear aperture of 22.9 mm and a thickness of 3 mm. The separation angle between the central ray of adjacent beamlets is 0.028^×^ 0.028 degrees. This generates a pattern with a full angle of 0.17^×^ 0.17 degrees. Both sides of the DOE are AR-coated for 405 nm, and the zero-order spot in the middle is specified to be 90-130% in intensity relative to the other spots, to ensure high uniformity. The width and timing of square pulses input to the modulation port on the laser power supply are controlled by an Agilent 3320A Waveform Generator. The upper objective used for photoactivation is a 60x Nikon water-immersion lens with NA = 1.00 and a working distance of 2 mm. A z-axis piezo holding this objective is used to finely adjust the image plane location of the beamlets produced by the DOE. The DOE photoactivation module is mounted on the condenser arm of an IX83 inverted microscope (Olympus) with a custom adaptor. The lower imaging objective (60x Olympus oil-immersion, NA = 1.35) is used to acquire epifluorescence images of photoactivated GFP. The fluorescence light source is an Olympus U-HGLGPS, which uses a 130 W mercury vapor short arc bulb and a fiber optic light guide to couple the source to the microscope. A GFP filter cube (470/40 EX; 525/5 EM; 495LP BS; set 49002, Chroma) separates excitation and emitted light. Images are recorded with a scientific CMOS camera (ORCA-FLASH 4.0 LT, Hamamatsu) with a 6.5 μm x 6.5 μm pixel size. At 60x magnification, the conversion factor between pixels and distance at the sample is 9.27 pixels/μm.

**Figure 1.**
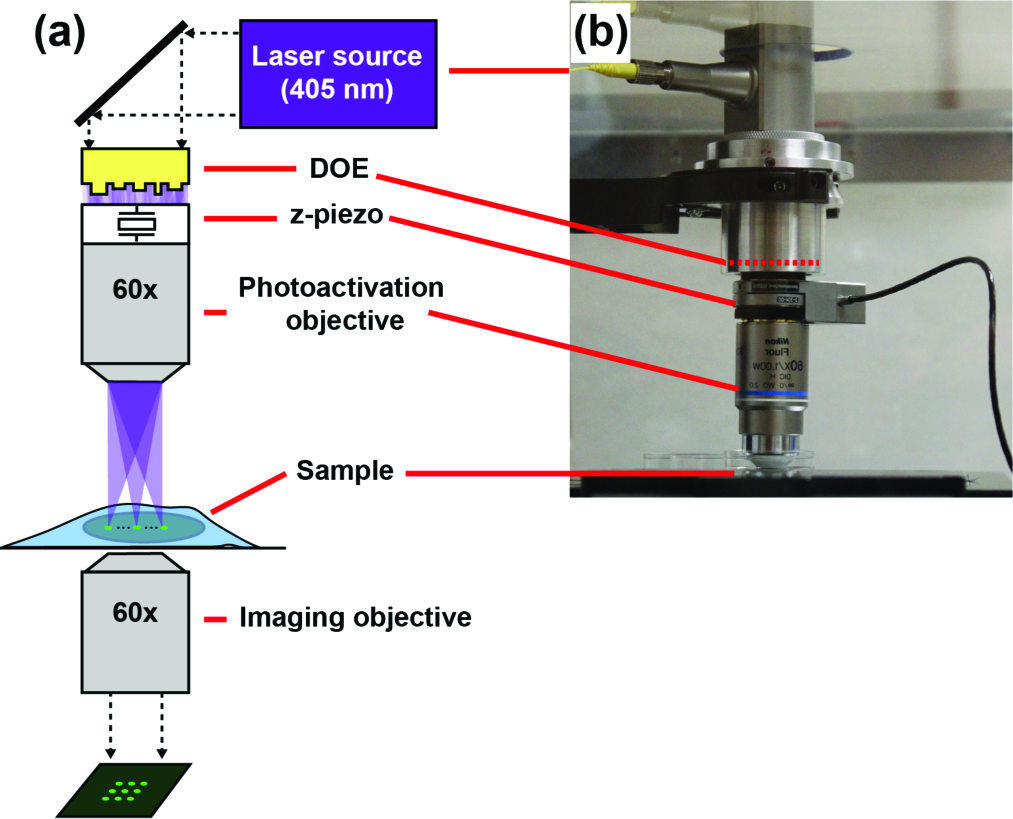
DOE photoactivation module. a) Schematic of the instrumentation. b) Photograph of the custom DOE module mounted on the condenser arm of an Olympus IX-83 inverted microscope.

### 2.2 System characterization

To characterize the temporal profile of the laser photoactivation pulse, we used a Thorlabs microscope slide power sensor (S170C) and energy meter console (PM100D and DET200). The analysis showed that when the power supply of the 405 nm laser diode was controlled with a 0.5 ms square pulse (of 250 mV amplitude), the time-course of the emitted light exhibited a shark's tooth pattern (Fig. 2a). The peak power for the whole pattern was approximately 12 mW at the highest setting of driving amplitude. To estimate the photon flux in one of the 49 DOE-generated spots, we multiplied the peak power of 7.6 mW measured at the sample for the whole array of beams by the efficiency (0.76), and divided by the number of spots (49) to obtain ~120 μW in a spot, corresponding to a photon rate of R = 1.8x10^10^ photons/s. Given a spot area of ~ 0.78 μm^2^, the photon flux F = R/A = 2.3x10^10^ photons/μm^2^s (I = 170 mW/mm^2^). This number is well below the photon flux of 1x10^12^ photons/μm^2^s, determined as a phototoxic threshold in eukaryotic cells (15).

**Figure 2.**
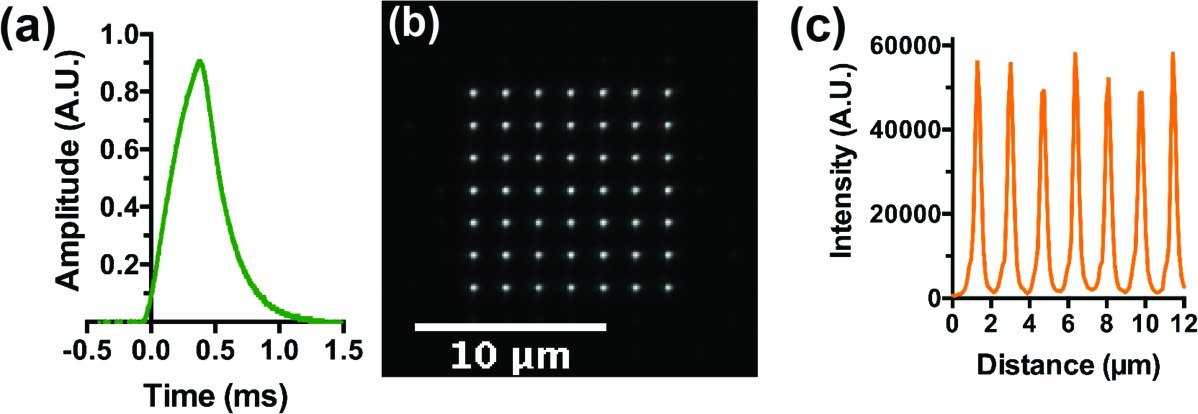
System characterization. a) Time profile of the laser pulse produced after driving the 405 nm laser modulation port with a 0.5 ms square pulse. b) Matrix of spots generated by the DOE module and directly visualized by bright field imaging, using only a #1.5 coverslip and water/oil immersion media in the light path. c) Intensity profile along one row of the spot matrix.

As an initial characterization of the spot array generated by the DOE module, images of the array were directly recorded using the brightfield setting of the microscope (Fig. 2b). This setup was ideal in that no optical medium other than distilled water was in the light path. Under these conditions, individual spots were 340±30 nm in diameter, defined as the full-width at half-maximum of intensity (FWHM; Fig. 2c).

### 2.3 Image Preprocessing

We used a photoactivatable fluorescent dye (rhodamine Q caged with ortho-nitroveratryloxycarbonyl; NVOC-RhQ; (16)) embedded in clear epoxy resin to test the DOE’s performance and to establish the image preprocessing method. After photoactivation with the 405 nm laser, an array of NVOC-RhQ spots was detected using a TxRed/rhodamine filter set (Fig. 3a).

**Figure 3.**
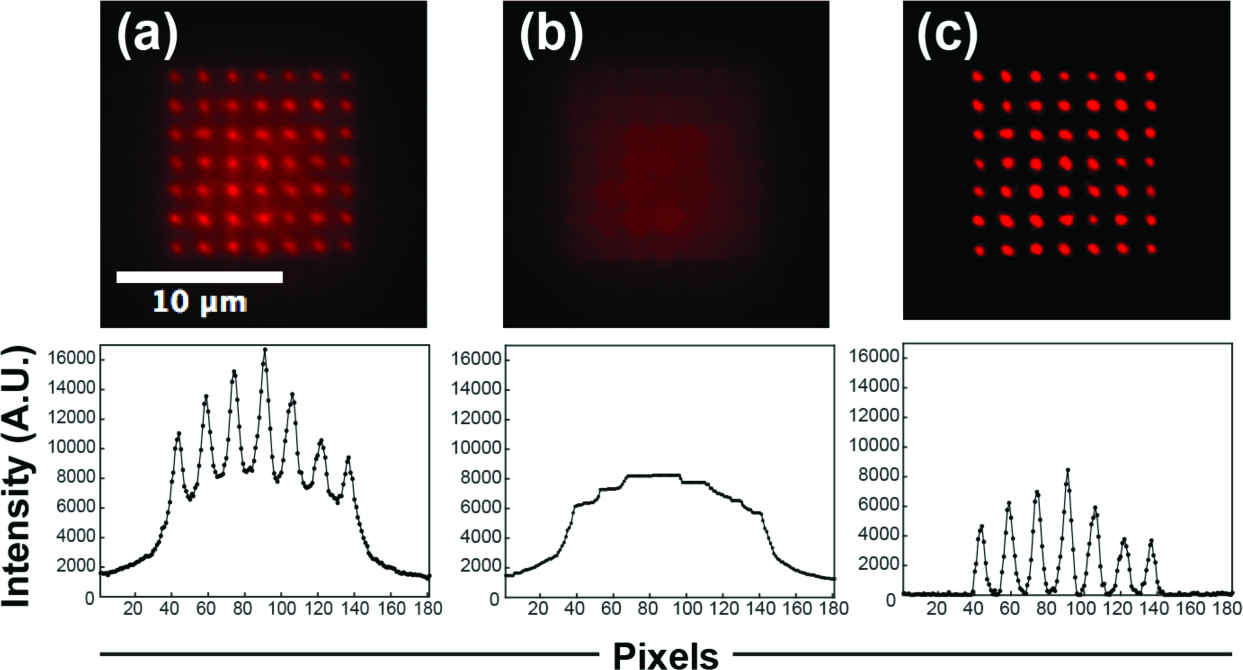
Image preprocessing. a) Fluorescence image generated after photoactivation of the NVOC-RhQ dye in clear epoxy resin. b) Background image (see text). c) Background-subtracted NVOC-RhQ photoactivation image. Intensity profiles are shown below each image. Each dot corresponds to one pixel (equivalent to 0.1 μm).

The intensity profile across the photoactivated NVOC-RhQ dot matrix revealed a “pedestal” of fluorescence between spots compared to direct imaging in water. This background is likely caused by the scattering of 405 nm light in the non-ideal epoxy medium, leading to NVOC-RhQ photoactivation under and between the intended spots. The amplitude and shape of this background pedestal were estimated by morphologically opening the raw image with a ball structuring element in Matlab (Fig 3b). The pedestal was subtracted from the original image, yielding the background-subtracted image shown in Fig 3c. The FWHM of photoactivated NVOC-RhQ spots was 640±60 nm (n = 5 frames; Fig 3c).

## 3 Results

### 3.1 Chromatin marker photoactivation within fixed and live cells

To assess the capabilities of the DOE photoactivation system in cell biology assays, we generated a stable osteosarcoma (U2OS) cell line expressing PAGFP fused to histone H2A (PAGFP-H2A). Cells were cultured on glass-bottom 35 mm dishes. First, laser intensity and pulse duration were varied to optimize PAGFP photoactivation while minimizing phototoxic damage (8). As shown in Fig. 4a, laser powers above 7.55 mW resulted in rapid photobleaching of PAGFP, as evidenced by the drop in PAGFP intensity after 20 and 30 ms of cumulative exposure to 9.94 and 11.70 mW peak laser powers, respectively. Line profiles of the dot matrices indicated that short photoactivation times (<10 ms) were needed to minimize photoactivation outside of the intended matrix. This undesirable photoactivation is likely caused by 405 nm light scattering. We anticipate that short photoactivation times have the added advantages of minimizing phototoxic effects. These results and the photodamage constraints indicated that 1 ms pulses of 7.55 mW laser power were most suitable for chromatin tracking experiments. With these settings, spot arrays generated in fixed PAGFP-H2A cells had a FWHM of 560±70 nm (Fig. 4b-c), which is 2.8 times the limit set by diffraction of PAGFP emission light (X/2NA = 200 nm). Similar FWHM values (630±100nm) were obtained with live cells (Fig. 4d).

**Figure 4.**
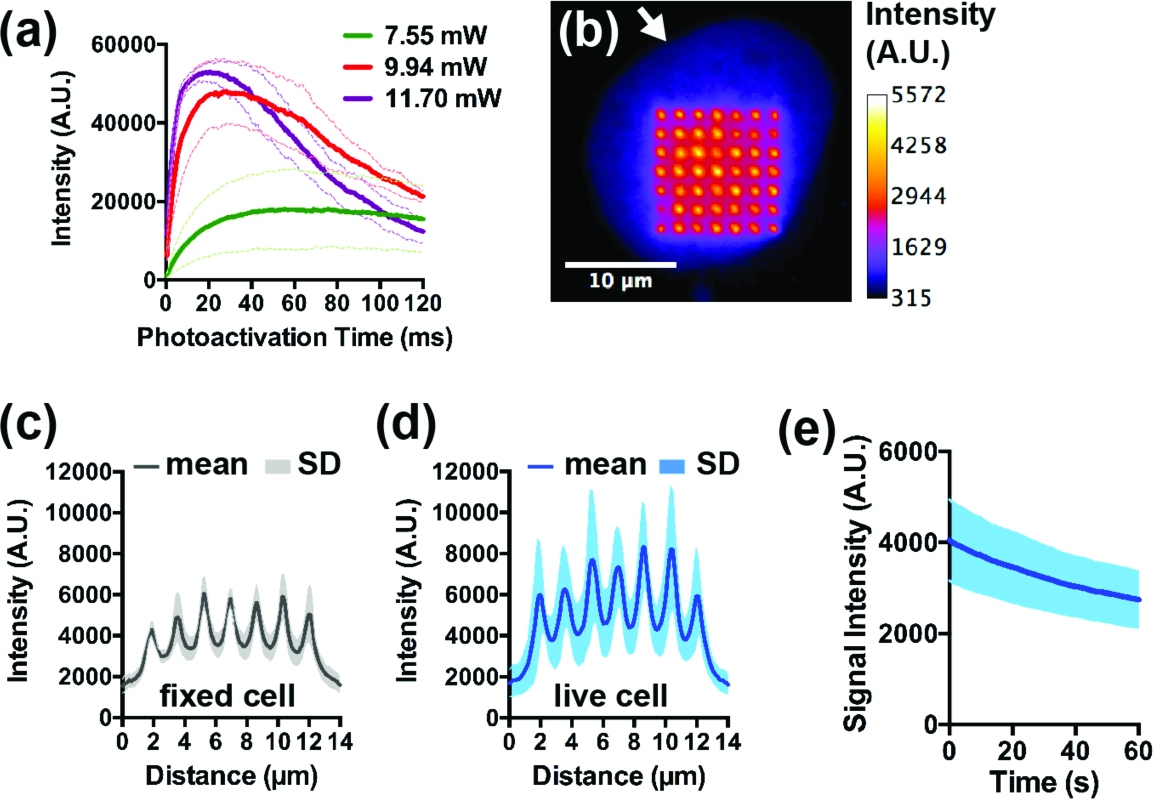
Chromatin marker photoactivation with DOE. a) GFP emission curves in fixed U2OS cell nuclei after photoactivation with three levels of 405 nm laser power. Nuclei were repeatedly photoactivated with 0.5 ms pulses with the total photoactivation time ranging from 0.5 – 120 ms. GFP fluorescence was imaged with a GFP filter cube. Average spot intensities and SD intervals (dark and pale lines, respectively; n = 3-4 cells) are plotted as a function of the laser exposure time. b) U2OS-PAGFP-H2A cell nucleus (arrow) after photoactivation (1 ms; 7.55 mW). Fluorescence intensity is visualized as a heat map. c-d) Profile plots of PAGFP in fixed cells (c) and in live cells (d) after photoactivation. e) Photoactivated GFP intensity in live cells as a function of GFP excitation time to show the bleaching rate.

To acquire time-lapse datasets for measurements of chromatin diffusion (*D*), the environmental chamber of the microscope was set to 37°C, and photoactivated PAGFP-H2A spots were imaged by epifluorescence microscopy in live cells for one minute at a 3.16 fps frame rate. Imaging conditions were optimized to minimize the bleaching rate (Fig. 4e). For comparison, cells were fixed with paraformaldehyde and imaged identically to live cells. Each dataset consisted of 200 frames. Representative time-lapse recordings of fixed and live cells (after registration - see below) are shown in Videos 1 and 2, respectively.

**Video 1.**
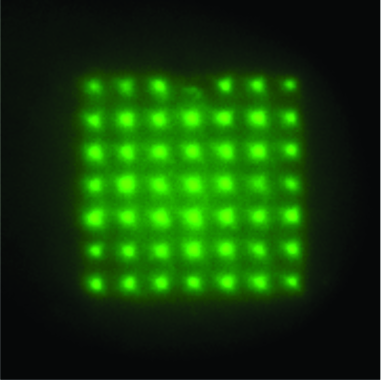
Time lapse imaging of a fixed U2OS cell expressing PAGFP-H2A (AVI, 674 KB).

**Video 2.**
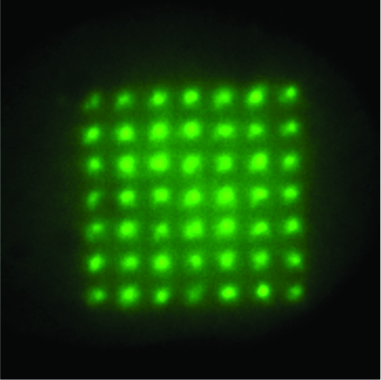
Time lapse imaging of a live U2OS-PAGFP-H2A cell (AVI, 679 KB).

### 3.2 Chromatin tracking

Cell migration as well as slight drifts of the microscope stage can confound measurements of chromatin motions. Hence, translations and rotations of the cell nucleus were first removed by sequential image registration using the StackReg plugin (17) for ImageJ. Next, using in-house Matlab code, background fluorescence was subtracted as in Fig 3. Importantly, this step removed photoactivated spot asymmetries, thereby improving Gaussian fits. Then, spots were tracked following the general approach of Crocker and Grier (18), but with modifications. Since the spots were in a known pattern, each image was divided into 49 regions of interest, each containing only one spot. This made data analysis more robust. The center of each spot was then localized to subpixel precision in each frame by fitting the spot intensity to a 2-dimensional Gaussian function (19). Finally, the diffusion coefficient of each spot was evaluated from the slope of the first 12 points in its mean-squared displacement curve (0.30-3.6s).

As expected, averaged *D* values were significantly higher (by a factor of 25), and more heterogeneous, in live compared to fixed cells (Fig. 5a-b; F test of variance, *P* < 0.0001). Importantly, values of *D* measured with the DOE system in live cells closely matched previous measurements of chromatin diffusion in mammalian cells performed with different methods (20).

**Figure 5.**
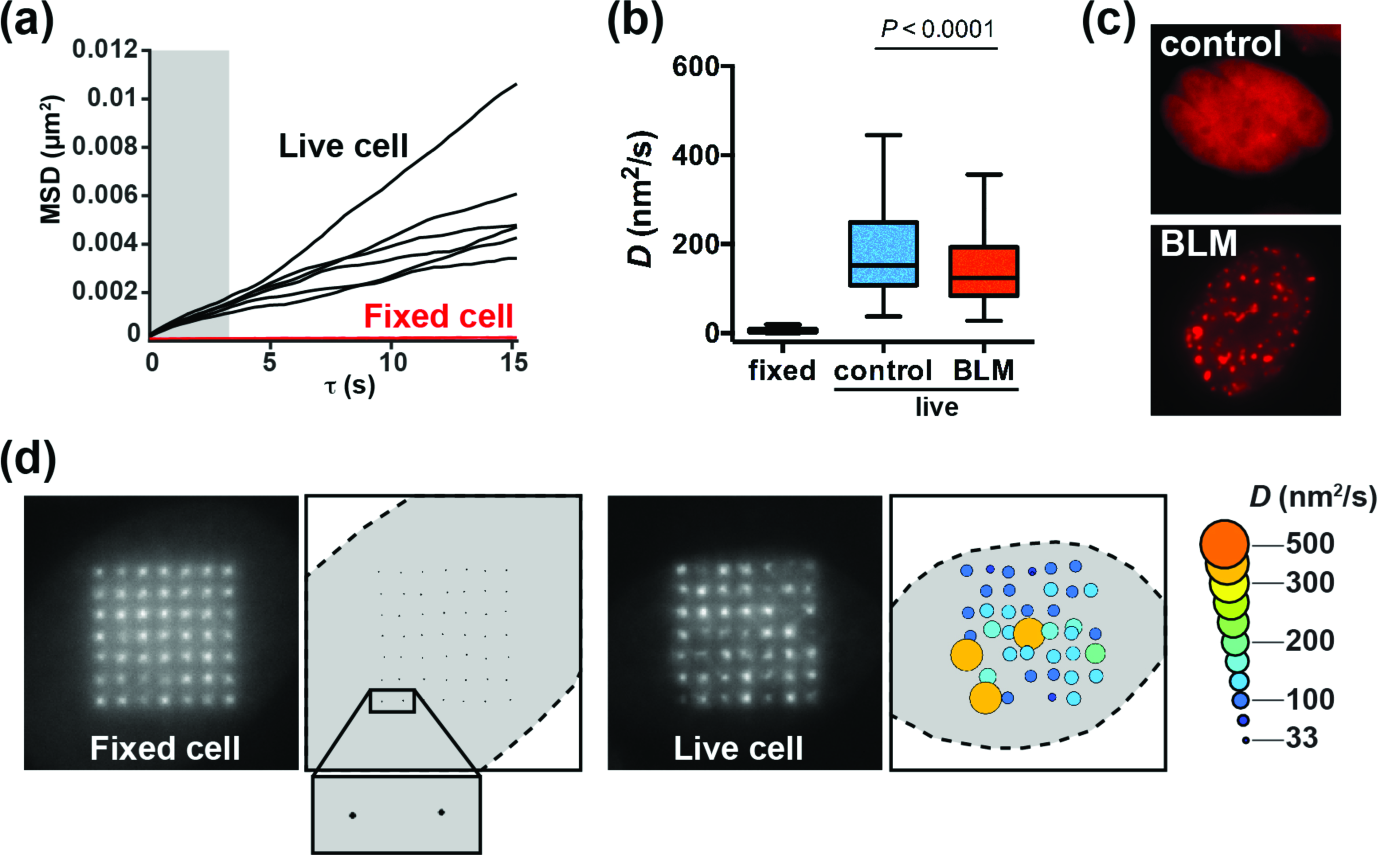
Mapping of chromatin motions in adherent cells. a) Representative MSD curves used to evaluate *D*. Data are shown for 6 spots in one live cell (black lines) and 6 spots in one fixed cell (red lines). *D* was obtained from the slope of the MSD, using the first 12 values of tau (grayed region of the graph). b) Box-and-whisker plot (Tukey method) of chromatin diffusion (D) measured in live and fixed cells. Live cells were either untreated (control) or treated with the chemotherapeutic drug bleomycin (BLM, 20mU/ml for 1h). c) Representative images of the DNA damage marker mCherry-53BP1ct in cells treated with BLM and in untreated cells. d) Maps of chromatin motions in fixed and live cells. The amplitude of *D* is visualized at specific nuclear regions with the size and colors of the circles. Small motions for fixed cells are visualized in the inset.

We reported previously that DNA damage causes a transient decrease in chromatin diffusion in U2OS cells (21). This effect was clearly apparent when comparing *D* values from cells treated with the radiomimetic drug bleomycin (BLM) to control, untreated cells (Fig. 5a). DNA damage induction by BLM was verified by expressing a DNA damage reporter (the C-terminal fragment of 53BP1 fused to mCherry; mCh-53BP1ct) (Fig. 5c). This marker was used to select cells for analysis. Spatial distributions of *D* values were visualized in fixed and live cells using bubble maps (Fig. 5d). The maps revealed unexpected levels of heterogeneity of chromatin diffusion at different locations in the cell nucleus. Follow-up studies will combine these maps with other spatial information of the cell nucleus (e.g., chromatin condensation, radial position, localization of nuclear bodies, etc.).

To determine if chromatin motions are coherent at the microscale level of our analyses, Pearson’s r correlation was computed for movements of all pairs of spots. Specifically, we were interested in whether changes in position from one frame to the next frame were correlated for the different spots. To quantify this we computed the differences in x‐ and y-coordinate (dx, dy) for adjacent time series frames for each spot, and then calculated the correlation coefficients of these 199 changes in x‐ and y-coordinate for all 49 spots (200 time frames of position data were the raw data used for this analysis). As a control, seven synthetic datasets (“cells”) were generated, each consisting of 200 frames with 49 spots. Independent Brownian motion was assigned to each spot, moving the spot from frame to frame with a realistic diffusion coefficient. Pearson’s r correlation was computed for dx and dy, considering each possible pair between the 49 spots in a “cell” (N=1176). Correlation values for the 7 “cells” were averaged and are plotted as heat maps in Fig. 6a. As expected, the synthetic data showed no correlation (r_dx_ = 0.0004±0.03 and r_dy_ = ‐0.00007±0.03; mean r ± SD). The same approach was applied to live cells. When considering all pairs of spots, no correlation was observed (rdx = ‐0.01±0.05; rdy = ‐0.01±0.05). However, some regions of the heat map stood out. Specifically, spots from opposite locations within the 7x7 grid (upper left corner) tended to be anti-correlated, whereas spots closer to one another (lower left and upper right) tended to be correlated. More strikingly, nearest neighbor (NN) positions were clearly more correlated than average (rdx = 0.08±0.07; rdy = 0.08±0.06; Fig. 6b).

**Figure 6.**
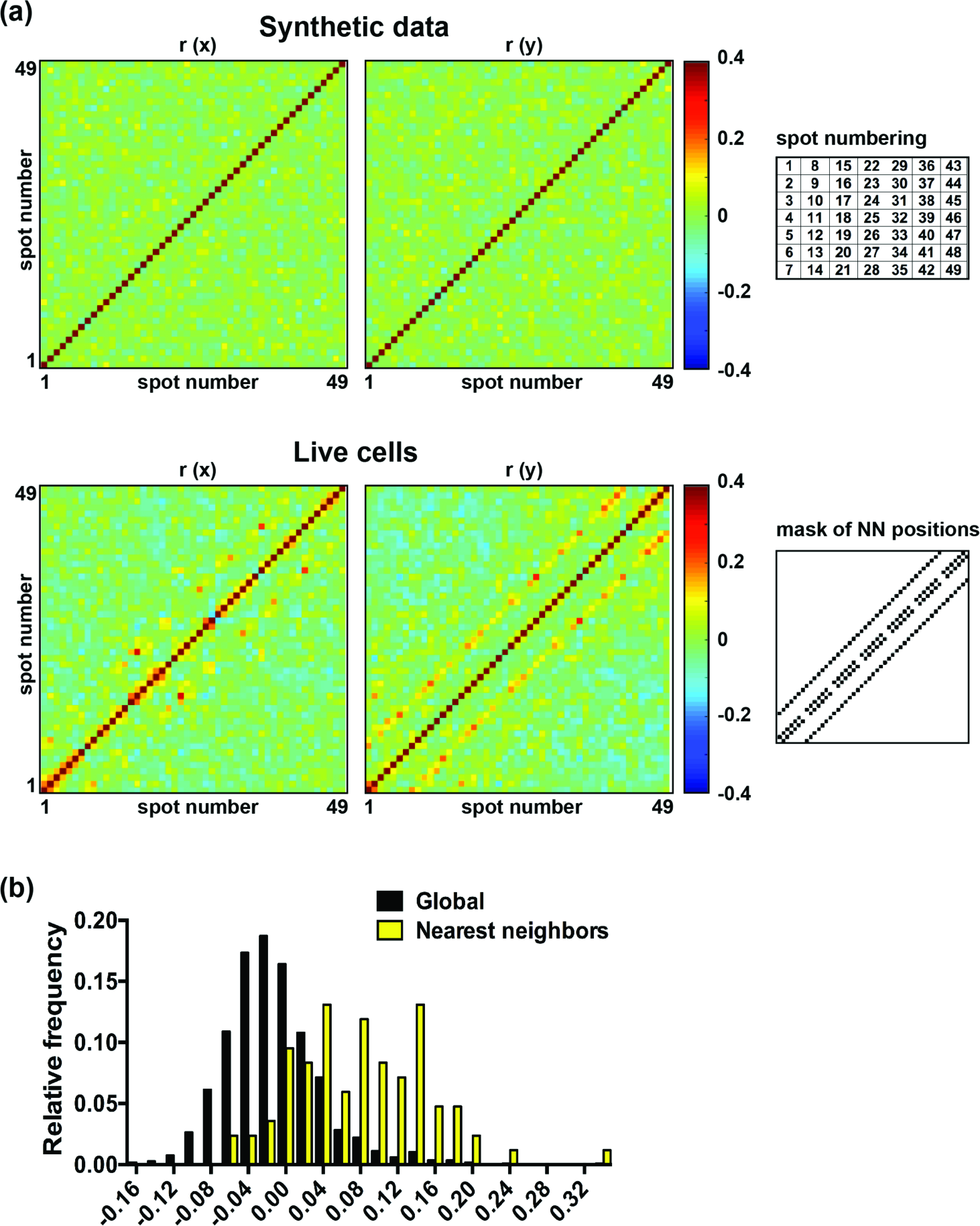
Correlation analysis of chromatin microdomain motions. a) Heat maps of correlation (Pearson’s r coefficients) for all pairs of photoactivated spots, in x (left) and y (right). Synthetic data were generated by assigning Brownian motions to arrays of 49 spots. The plots represent averages (N=7). The positions of the nearest neighbors (NN) within the plots are shown on the right of the middle figures. To the right of the upper correlation plots we have provided the numbering scheme of spots used to generate the correlation maps. b) Probability distributions of r values computed from live cells and corresponding to changes in x between frames (motions along x).

Chromatin motions are necessary for the biogenesis of genomic translocations (22) that drive hematologic malignancies. We therefore anticipate applications of the method presented herein for the study of chromatin dynamics in the context of the hematopoietic system, and developed a protocol for analysis of (non-adherent) CD34+ hematopoietic stem/progenitor cells (HSPCs) (Fig. 7). Briefly, fresh HSPC were maintained for 24h in culture, then nucleofected with PAGFP-H2A and mCherry plasmids using an Amaxa protocol (Lonza). 30h later cells were immobilized on glass coverslips using a cell adhesive (CellTak, Corning) and embedded in a hydrogel topped with HSPC medium. Nucleofected cells are easily identified based on mCherry fluorescence, then PAGFP-H2A is photoactivated with the DOE system and time-lapse images recorded, as for the adherent U2OS cells.

**Figure 7.**
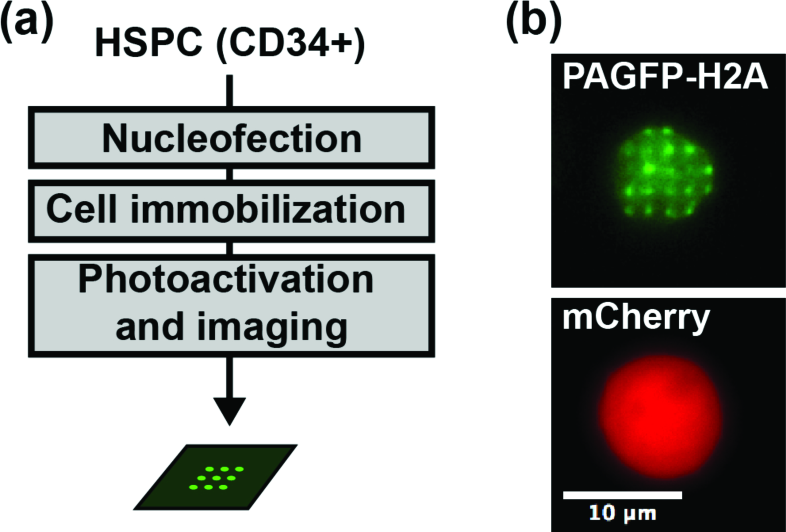
Chromatin motions in nonadherent hematopoietic stem/progenitor cells (HSPCs). a) Flow chart of the procedure for sample preparation and data acquisition. b) DOE photoactivation of PAGFP-H2A in HSPCs, giving ~20 bright spots on a green background. Nucleofected cells were identified based on red fluorescent signals from mCherry, which was co-expressed with PAGFP-H2A.

## 4 Discussion

We have developed a facile approach based on structured illumination to quantify the movement of photoactivated microdomains of the cell with nanometer accuracy. One application of this method is illustrated by tracking chromatin motions. Based on spot sizes, the number of nucleosomes tracked collectively is estimated at 10^5^. Therefore, the method yields mesoscale readouts of global chromatin motions at specific sub-domains of the cell nucleus, which are complementary to other tracking approaches where shorter stretches of chromatin are labeled. Beyond measurements of chromatin motions, we anticipate a broad application range of the method to measure diffusion, with the advantage of reducing phototoxic damage to cells compared to photobleaching-based techniques (23). Our method can simultaneously map distinct spatial regions, unlike FRAP and Raster Imaging Correlation Spectroscopy (24).

Previously, a paired-particle tracking approach was implemented to determine chromatin *D*, using a confocal microscope for photoactivation and imaging of two PAGFP-H2A spots (21). With this pairwise method, or with a more recent extension of the confocal approach to 9 spots (unpublished results), photoactivated spots could be tracked up to one minute (at 3.3 fps). Then, photobleaching prevented tracking. The new DOE-based design has the following advantages over these previous approaches. First, the intensity pattern is static and defined. This reproducibility greatly eases analyses. Second, the method is simple. It involves a passive optical element that requires no adjustment. Third, it is well-modeled, since each photoactivation beamlet has a known, quasi-identical intensity. Hence, intensity patterns detected in cells represent the expression level of the reporter in different cellular regions rather than illumination artifacts. The fourth advantage is speed. All spots are illuminated simultaneously. The fifth advantage is ease of integration. The DOE module can simply be mounted onto existing microscope platforms. Sixth, the current approach is flexible. Spot patterns can be changed by swapping one DOE for another. The seventh advantage is the high quality patterns produced by the DOE, which result in narrow spots (~ 40% smaller than those achieved with a Zeiss CLSM710 confocal system). The final (eighth) advantage is the high spatial sampling capability of the DOE method, due to the added speed and spot resolution. We estimate a limit of sampling density of 1 μm based on calculated FWHM values, corresponding to 30-300 spots/nucleus depending on cell types, although this limit may eventually be limited by the extent of the chromatin motion. One limitation of the method is the possibility of vibrations from acoustical or mechanical sources causing pattern blur or distortion, in particular when the DOE module is mounted on a condenser mount. Vibration effects can be minimized to being negligible by quick laser pulsing and were found not to be an issue for short photoactivation times. A second challenge is that a small amount of scattered light produces background activation of the reporter that needs to be subtracted post-imaging for optimal tracking results.

## 5 Conclusions

Maps of chromatin motions in live cells have been obtained by a new method based on diffractive optics and photoactivatable chromatin reporters. The maps hint that correlated motions exist between adjacent focal volumes in chromatin microdomains at the time scale used for analysis. The imaging approach may lead to a better understanding of the mechanisms regulating chromatin dynamics in normal and pathological contexts. Application of this method to clinical samples will allow us to test the hypothesis that chromatin motions predict myeloid neoplasms caused by genomic translocations.

## Acknowledgments

We thank Bob Morris for assistance with precision machining for the custom optical setup and Dr. Jacques Neefjes (the Netherland Cancer Institute) for providing the PAGFP-H2A DNA construct and Dr. T. de Lange (the Rockefeller University) for providing the mCherry-53BP1ct DNA construct. This work was funded by the National Cancer Institute (R00CA163957); National Cancer Institute’s Cancer Center Support Grant award number P30CA012197 issued to the Wake Forest Baptist Comprehensive Cancer Center; Wake Forest Clinical and Translational Science Institute (CTSI) Open Pilot grant; the Wake Forest Center for Molecular Signaling (CMS) via its imaging facility. Research reported in this publication is solely the responsibility of the authors and does not necessarily represent the official views of the National Cancer Institute.

**Keith Bonin** is a Professor of Physics at Wake Forest University. He received his BS degree in physics from Loyola University in New Orleans in 1978, and his PhD degree in physics from the University of Maryland in 1984. He is the author of more than 60 journal articles and one book. His current research interests include optics, biophysics, microscopy, and biophotonics. He is a member of SPIE.

**George Holzwarth** is a Research Professor of Physics at Wake Forest University. He received his BA in physics from Wesleyan University in 1959 and his PhD in Biophysics from Harvard University in 1964. He is the author of more than 70 journal articles and holds 3 patents. His current research interests are molecular motor mechanics, intracellular microrheology, and the motions of chromatin within cellular nuclei. Primary tools are DIC and fluorescence video microscopy.

**Pierre-Alexandre Vidi** is an Assistant Professor of Cancer Biology at Wake Forest University Health Sciences. He received his MS degree in biology from the University of Lausanne, Switzerland, in 2002 and his PhD degree in plant physiology from the University of Neuchâtel, Switzerland, in 2006. He has authored 20 peer-reviewed journal publications as well as seven reviews and book chapters. Research in his newly established laboratory focuses on the cellular responses to DNA damage.

Biographies for the other authors are not available.

## References

1. V. Dion, and S. M. Gasser, “Chromatin movement in the maintenance of genome stability,” Cell 152(6), 1355-1364 (2013).

2. W. F. Marshall et al., “Interphase chromosomes undergo constrained diffusional motion in living cells,” Current biology: CB 7(12), 930-939 (1997).

3. V. Dion et al., “Increased mobility of double-strand breaks requires Mec1, Rad9 and the homologous recombination machinery,” Nature cell biology 14(5), 502-509 (2012).

4. J. Mine-Hattab, and R. Rothstein, “Increased chromosome mobility facilitates homology search during recombination,” Nature cell biology 14(5), 510-517 (2012).

5. H. Ma et al., “Multicolor CRISPR labeling of chromosomal loci in human cells,” Proceedings of the National Academy of Sciences of the United States of America 112(10), 3002-3007 (2015).

6. B. Chen et al., “Expanding the CRISPR imaging toolset with Staphylococcus aureus Cas9 for simultaneous imaging of multiple genomic loci,” Nucleic acids research 44(8), e75 (2016).

7. A. Jacome, and O. Fernandez-Capetillo, “Lac operator repeats generate a traceable fragile site in mammalian cells,” EMBO reports 12(10), 1032-1038 (2011).

8. G. H. Patterson, and J. Lippincott-Schwartz, “Selective photolabeling of proteins using photoactivatable GFP,” Methods 32(4), 445-450 (2004).

9. M. J. Kruhlak et al., “Changes in chromatin structure and mobility in living cells at sites of DNA double-strand breaks,” The Journal of cell biology 172(6), 823-834 (2006).

10. K. Wiesmeijer et al., “Chromatin movement visualized with photoactivable GFP-labeled histone H4,” Differentiation; research in biological diversity 76(1), 83-90 (2008).

11. M. Saxena, G. Eluru, and S. S. Gorthi, “Structured illumination microscopy,” Adv. Opt. Photon. 7(2), 241-275 (2015).

12. M. F. Langhorst, J. Schaffer, and B. Goetze, “Structure brings clarity: structured illumination microscopy in cell biology,” Biotechnol J 4(6), 858-865 (2009).

13. T. Lukeš et al., “Comparison of image reconstruction methods for structured illumination microscopy,” Proc. of SPIE 9129(91293J), (2014).

14. D. L. Andrews, Structured Light and Its Applications: An Introduction to Phase-Structured Beams and Nanoscale Optical Forces, Academic Press, Burlington, MA (2008).

15. P. M. Carlton et al., “Fast live simultaneous multiwavelength four-dimensional optical microscopy,” Proceedings of the National Academy of Sciences of the United States of America 107(37), 16016-16022 (2010).

16. L. M. Wysocki et al., “Facile and general synthesis of photoactivatable xanthene dyes,” Angewandte Chemie 50(47), 11206-11209 (2011).

17. P. Thevenaz, U. E. Ruttimann, and M. Unser, “A pyramid approach to subpixel registration based on intensity,” IEEE Trans Image Process 7(1), 27-41 (1998).

18. J. C. Crocker, and D. G. Grier, “Methods of Digital Video Microscopy for Colloidal Studies,” Journal of Colloid and Interface Science 179(298-310 (1996).

19. A. Yildiz et al., “Myosin V walks hand-over-hand: single fluorophore imaging with 1.5nm localization,” Science 300(5628), 2061-2065 (2003).

20. J. Mine-Hattab, and R. Rothstein, “DNA in motion during double-strand break repair,” Trends Cell Biol 23(11), 529-536 (2013).

21. J. Liu et al., “Nanoscale histone localization in live cells reveals reduced chromatin mobility in response to DNA damage,” Journal of cell science 128(3), 599-604 (2015).

22. V. Roukos et al., “Spatial dynamics of chromosome translocations in living cells,” Science 341(6146), 660-664 (2013).

23. J. W. Dobrucki, D. Feret, and A. Noatynska, “Scattering of exciting light by live cells in fluorescence confocal imaging: phototoxic effects and relevance for FRAP studies,” Biophysical journal 93(5), 1778-1786 (2007).

24. M. J. Rossow et al., “Raster image correlation spectroscopy in live cells,” Nature protocols 5(11), 1761-1774 (2010).

